# Cell morphological motif detector for high-resolution 3D microscopy images

**DOI:** 10.1101/376608

**Authors:** Meghan K. Driscoll, Erik S. Welf, Kevin M. Dean, Reto Fiolka, Gaudenz Danuser

## Abstract

Recent advances in light-sheet microscopy enable imaging of cell morphology and signaling with unprecedented detail. However, the analytical tools to systematically measure and visualize the intricate relations between cell morphodynamics, intracellular signaling, and cytoskeletal dynamics have been largely missing. Here, we introduce a set of computer vision and graphics methods to dissect molecular mechanisms underlying 3D cell morphogenesis and to test whether morphogenesis itself affects intracellular signaling. We demonstrate a machine learning based generic morphological motif detector that automatically finds lamellipodia, filopodia, and blebs on various cell types. Combining motif detection with molecular localization, we measure the differential association of PIP_2_ and Kras^V12^ with blebs. Both signals associate with bleb edges, as expected for membrane-localized proteins, but only PIP_2_ is enhanced on blebs. This suggests that local morphological cues differentially organize and activate sub-cellular signaling processes. Overall, our computational workflow enables the objective, automated analysis of the 3D coupling of morphodynamics with cytoskeletal dynamics and intracellular signaling.

## Introduction

Cellular morphogenesis is critical to single and collective cell migration,^1,2^ tissue homeostasis and remodeling,^3^ environmental sensing,^4^ and cell-cell communication.^5^ Morphogenesis is driven by forces generated primarily by the assembly and contraction of the actomyosin cytoskeleton.^6^ These mechanical processes are downstream of chemical signaling activities. The cascade from signaling to cytoskeleton dynamics and finally to cell morphology has been extensively studied. For example, the formation of the three common morphological structures, lamellipodia, blebs, and filopodia (Fig. 1a-c and Supplementary Fig. 1), depends on well-characterized assemblies of actin filaments (Fig. 1d-f).^7^ How morphology, in turn, may govern signaling has been little investigated. Morphology likely participates as a key element in signal transduction pathways via mechanisms such as preferential protein interaction with membranes of particular curvature,^8^ or modulation of the concentration and diffusion of signaling components,^9,10^ which affects the underlying biochemical reaction cascade.

**Figure 1.**
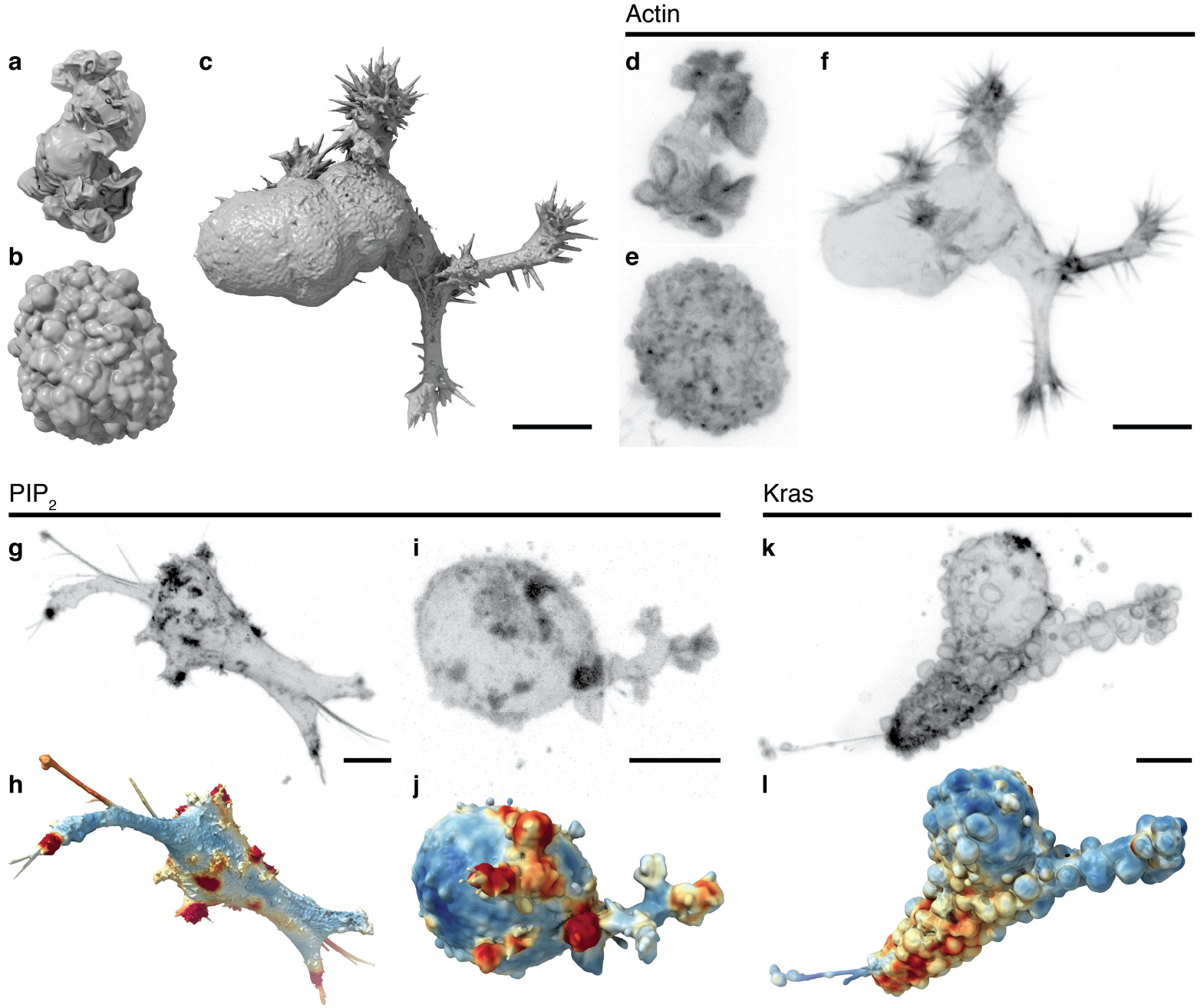
Cell morphology and signaling are coupled. Surface renderings of (**a**) a dendritic cell expressing Lifeact-GFP, (**b**) an MV3 melanoma cell expressing tractin-GFP, and (**c**) a human bronchial epithelial cell (HBEC) expressing tractin-GFP. (**d-f**) Maximum intensity projections (MIPs) of the cells shown in **a-c**, using an inverse look up table. Panels **a-f** are shown at the same scale. Additional views of these cells are shown in Supplementary Fig. 1. (**g**) A MIP of a branched MV3 cells expressing PLCΔ-PH-GFP, a PIP_2_ translocation biosensor. (**h**) A surface rendering of the same cell. Surface regions with relatively high PIP_2_ localization are shown in red, whereas regions of relatively low localization are shown in blue. (**i**) A MIP and (**j**) a surface rendering of a blebbing MV3 cell expressing PLCΔ-PH-GFP. (**k**) A MIP of an MV3 cell expressing GFP-Kras^V12^. (**l**) A surface rendering of **k**. Surface regions with relatively high Kras localization are shown in red, whereas regions of relatively low localization are shown in blue. (**m**) The morphological motif detection framework introduced here. Scale bars, 10 μm.

The integrated study of signaling and morphology at subcellular length scales has become possible with the recent advent of high-resolution 3D light-sheet microscopy.^11-16^ Critically, some of these microscopes allow observation of subcellular processes in cells that are deeply embedded in 3D tissue models where the morphology is not constrained by adhesion to a coverslip. To probe the putative relationships between signaling and morphology, we used microenvironmental selective plane illumination microscopy^15^ (meSPIM) to image the sub-cellular activity of two prototypical signaling molecules, PIP_2_ and transformed Kras^V12^, in cells embedded in 3D collagen. PIP_2_ is a membrane-bound phosphoinositide that plays critical roles in many signaling pathways.^17^ We unexpectedly found that PIP_2_ forms activation clusters in both branched (Fig. 1g,h) and blebbed cells (Fig. 1i,j). Three-dimensional renderings of the activation levels suggest that these clusters tend to colocalize with filopodial tufts (Fig. 1h) and blebs (Fig. 1j). Kras^V12^ is constitutively active with broad oncogenic functionality.^18^ Like PIP_2_, it appears to colocalize with certain morphological structures (Fig. 1k,l). These associations pose the question of whether rugged surface geometries generally associate with elevated signaling, and whether there are differences in the modes of association between PIP_2_ and Kras.

Answering such questions with statistical robustness requires the interpretation of massive quantities of 3D images using automated workflows. However, few computational tools are available to analyze these data.^19^ Here, we introduce computational tools combining approaches from computer vision, machine learning, and computer graphics to unravel the coupling between morphology, the cytoskeleton, and signaling. Since morphological structures, such as lamellipodia, blebs, and filopodia, tend to be associated with distinct signaling and cytoskeletal hierarchies, we developed a generic 3D *morphological motif* detector that identifies such structures. Combining the detector with computational tools to measure fluorescence localization near the cell surface, motif geometry, and boundary motion, in a proof-of-concept study we analyzed the association of Kras^V12^ and PIP_2_ with blebs. In the future, these methods will be instrumental to furthering our understanding of the feedback interactions between signaling, the cytoskeleton, and morphological dynamics in 3D.

## Results

### Detecting cellular morphological motifs

To detect morphological motifs, we extract the cell surface, decompose the surface into convex patches, optionally merge these patches, and then classify the patches by morphological motif (Fig. 2a-e). The cell surface is represented as a triangle mesh, which is the standard approach used in computer graphics to describe free form 3D objects. This representation enables the use of convenient mathematical formalisms for geometric analysis and the visualization of complex shapes. For most cells, we automatically extract the surface as an isosurface of the deconvolved 3D image (Fig. 2f-h). However, surface extraction is not trivial and sometimes requires that cell surface extraction parameters be tailored to the cell type and fluorescence label. For example, we extract the surfaces of actin-labeled dendritic cells as an isosurface of an image that combines the deconvolved image, an image with enhanced planar features, and an image with an enhanced cell interior, which has little actin (Supplementary Fig. 2). Although the cell surface extraction is dependent on the experimental system, the remainder of the workflow does not require customization and the same set of parameters were used throughout.

**Figure 2.**
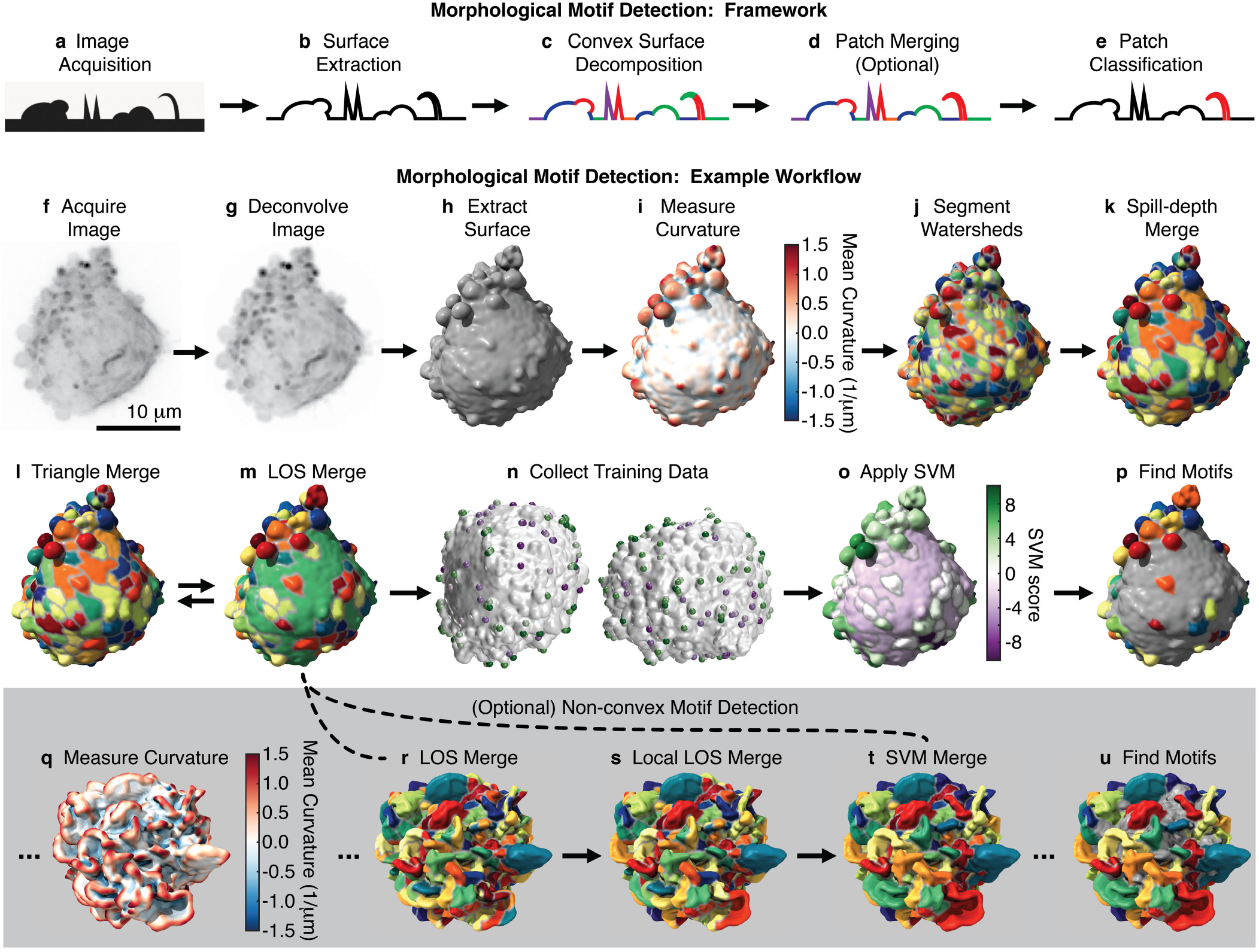
Morphological motif detection framework and example workflow. To detect morphological motifs, (**a**) following image acquisition, (**b**) we extract the cell surface, (**c**) decompose that surface into convex patches, (**d**) optionally merge those patches, (**e**) and then finally classify the patches by morphological motif. Panels **f-p** show our detection framework applied to a blebbed cell. (**f**) MIP of a 3D image of an MV3 melanoma cell expressing tractin-GFP. (**g**) MIP of the deconvolved image of the same cell. (**h**) The surface of the cell extracted from the deconvolved image as a triangle mesh. (**i**) The mean surface curvature of the cell. Regions of large positive curvature are shown red, flat regions are shown white, and regions of large negative curvature are shown blue. (**j**) A watershed segmentation of mean surface curvature. Segmented patches are shown in different colors. (**k**) A spill-depth based merging of the segmented patches. (**l**) A triangle-rule based merging of the patches. (**m**) A line-of-sight (LOS) based merging of the patches. The triangle and LOS rules are applied iteratively. (**n**) User generated training data for two different cells. Patches identified as “certainly a bleb” are shown green, whereas patches identified as “certainly not a bleb” are shown purple. (**o**) A support vector machine (SVM) classifier trained on user data applied to the cell. Patches shown in green have high SVM scores and a high inferred likelihood of being a bleb, whereas patches shown in purple have low SVM scores and a low inferred likelihood. (**p**) Detected blebs are shown randomly colored and non-blebs are shown gray. To detect non-convex motifs, such as lamellipodia, convex patches are merged as shown in **q-u**. (**q**) The mean surface curvature of a lamellipodial dendritic cell expressing Lifeact-GFP. (**r**) Convex surface patches for the same cell. (**s**) A local-LOS based merging of these patches. (**t**) An SVM based merging of the patches. The SVM was trained on user-supplied examples of adjacent patches that should certainly be merged and adjacent patches that should certainly not be merged. (**u**) Detected lamellipodia are shown randomly colored and non-lamellipodia are shown gray. Supplementary Fig. 4 shows an additional example of the bleb detection workflow.

We next decompose the surface into patches. Although various mesh segmentation algorithms have been developed for this purpose in computer graphics,^20^ they do not segment out biologically relevant features, and the benchmark data sets they were tested on lack the multitude of protrusion types common to cells imaged at high resolution.^21^ In our workflow, we first decompose the surface into convex patches and then merge these patches into convex regions that are locally as large as possible. We decompose the surface into convex patches since most protrusions are either convex, e.g. filopodia and blebs, or composed of multiple convex regions, e.g. lamellipodia and flagella. Indeed, by visual inspection people tend to partition 3D surfaces into convex regions,^22^ suggesting that canonical protrusions are likely convex or composed of multiple convex regions.

Decomposing a mesh into convex regions is an NP-complete problem,^23^ and thus is computationally intractable for large meshes, even with extensive computing resources. We therefore combine several techniques to segment the surface into approximately convex patches. First, we calculate the mean curvature at every face on the mesh, and then break the surface into small patches via a watershed-based segmentation of mean curvature (Fig. 2i,j).^24^ These small patches are computationally manipulated more easily than individual faces and are analogous to superpixels in image segmentation. Next, we merge adjacent patches using a spill-depth criterion.^24^ In this context, the spill-depth is the difference between the maximum curvature at the patch-patch interface and the minimum curvature inside the patches(Fig. 2k). The effect of varying patch-merging parameters, such as the spill-depth threshold, is shown in Supplementary Fig. 3.

Following spill-depth merging, we further iteratively merge patches using the line-of-sight (LOS)^25^ and the triangle^15^ criteria (Fig. 2l,m). The LOS criterion merges patches if the percentage of rays that connect the two patches without exiting the cell is above a certain threshold. Hence, fulfilling this criterion requires only approximate convexity between the patches. When left unconstrained, this criterion would merge a small patch into an adjacent large patch if the majority of the large patch extended to the far side of the cell. To overcome this problem, we only consider rays of length less than twice the smaller patch size. Similar to the law of cosines in trigonometry, the triangle criterion merges adjacent patches whose joint closure surface area is small compared to the sum of their individual closure surface areas. We define the closure surface area as the additional surface area needed to close the mesh composing the patch. The triangle criterion embodies the short-cut rule,^26^ which states that given multiple convex shape decompositions, the decomposition with the shortest cuts between segments is preferred. Collectively, the three patch-merging criteria decompose the cell surface into convex patches.

Convex patches are then classified by morphological motif type using a Support Vector Machine (SVM). For each patch, 23 geometric features are calculated, including standard geometric measures, such as perimeter and mean Gaussian curvature, and measures that have previously been developed for mesh segmentation, such as the shape diameter function^27^ (Supplementary Table 1). Features are automatically selected for each set of training data by successively removing randomly chosen features until prediction quality is hampered. Following SVM training (Fig. 2n) and feature selection, an SVM is used to classify patches by motif type (Fig. 2o,p).

The outcomes of machine learning approaches, such as SVMs, are critically dependent on training data quality. To generate training data for the SVM, we built an interface where users can rotate 3D surfaces, zoom in and out, and click on patches to identify them as protrusions. Presented with the same four randomly chosen cells and asked to identify blebs, three users chose 46±6% of patches. On the other hand, when asked to identify patches that were not blebs, the same three users chose 25±4% of the patches, hence classifying 75% of the patches as blebs. This discrepancy carried over into SVM models, where for the two training sets 45±7% and 77±6% of the patches were identified as blebs. Asking users to click only on patches that are certainly blebs and then only on patches that are certainly not blebs resulted in models that classified an intermediate percentage of patches, 52±6%, as blebs. On average, users classified 20±3% of the patches as certainly a bleb or certainly not a bleb. To avoid bias towards a particular protrusion morphology, we therefore train SVMs with data where users only choose patches they can confidently classify.

Although many morphological motifs, including blebs and filopodia, are described by a single convex surface patch, some motifs, such as lamellipodia, are composites of multiple convex patches. To detect these motifs, we merge convex patches prior to patch classification. Convex patches are first merged using a modified LOS criterion in which only short rays starting and ending near the patch-patch interface are considered (Fig. 2q-s). This criterion effectively merges adjacent patches with smoothly varying curvature along their interface. Second, we merge adjacent patches using a machine-learning framework. Thirty-six geometric features are calculated for each pair of patches (Supplementary Table 2). To generate training data, users click on adjacent patches that should certainly be merged and adjacent patches that should certainly not be merged. Following sequential feature selection, an SVM is used to merge patches (Fig. 2t). Merged patches are then classified by motif type, as described in the previous paragraph (Fig. 2u).

We trained models to detect blebs, filopodia, and lamellipodia (Fig. 3 and Supplementary Fig. 5). Importantly, the algorithm selected different geometric features to detect the three morphologies. To determine which geometric features best distinguished morphologies, starting from no features, we successively added the most discriminative feature to the model (Supplementary Table 3). The features that best distinguished blebs from non-blebs were volume / (closure surface area)^3/2^ and mean curvature on the protrusion edge. Closure surface area is the minimum amount of additional surface area needed to create a closed polygon from the mesh of the patch. The features that best distinguished filopodia from non-filopodia were the distance from the center of the closure surface area to the mean face position, a measure of morphological feature length, and patch surface area. This same measure of morphological feature length as well as patch volume were the best features for distinguishing lamellipodia from non-lamellipodia.

**Figure 3.**
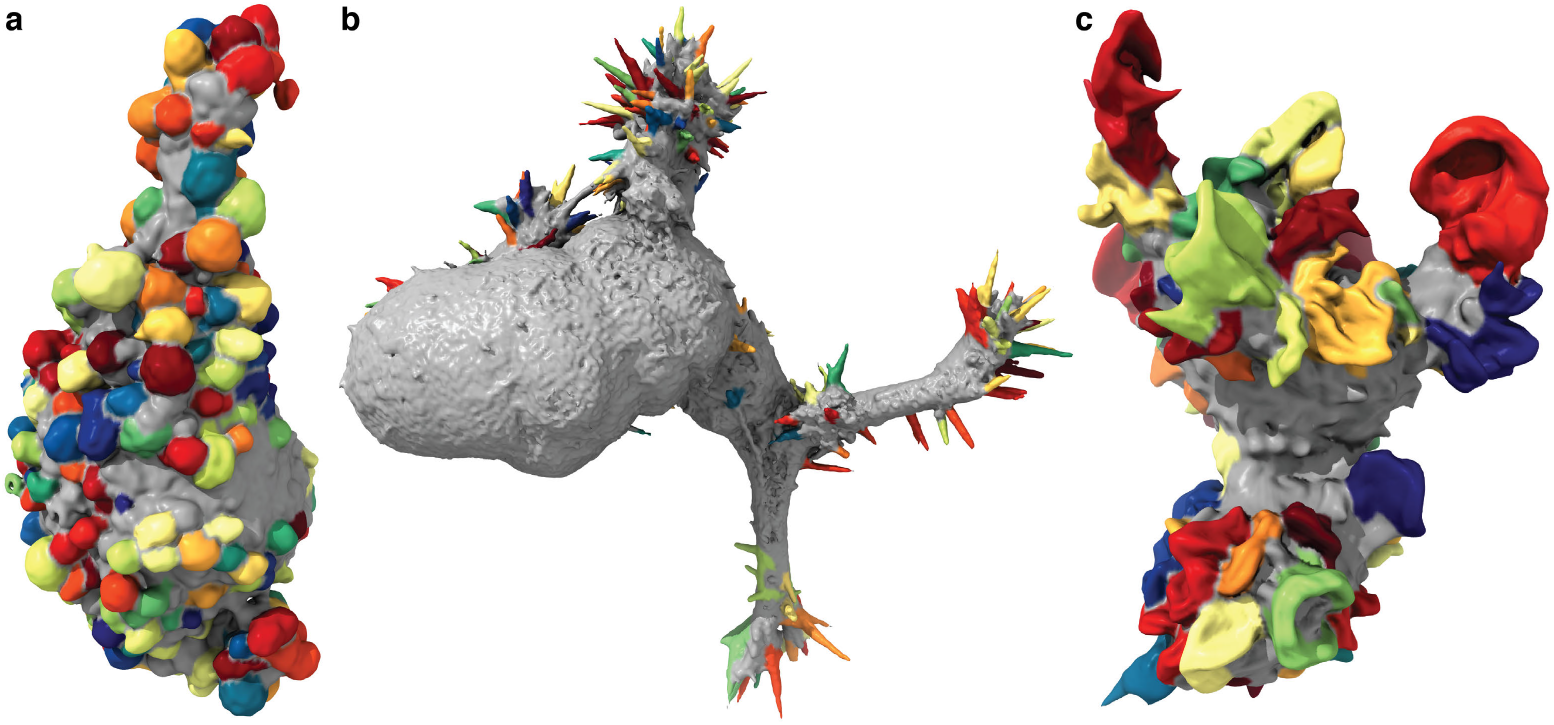
Detected blebs, filopodia, and lamellipodia. (**a**) Blebs detected on an MV3 melanoma cell, (**b**) filopodia detected on an HBEC cell, and (**c**) lamellipodia detected on a dendritic cell. Additional example detections are shown in Supplementary Fig. 5.

Most cells in our diverse data set showed predominately one protrusion type. However, as a proof-of-concept, we built a multiclass detector using a collection of melanoma cells that exhibited extensive blebs and small numbers of filopodia (Fig. 4a). To do so, we generated multiple SVM models in a one-vs-one framework in which separate models were used to distinguish each pair of morphological structure types.

**Figure 4.**
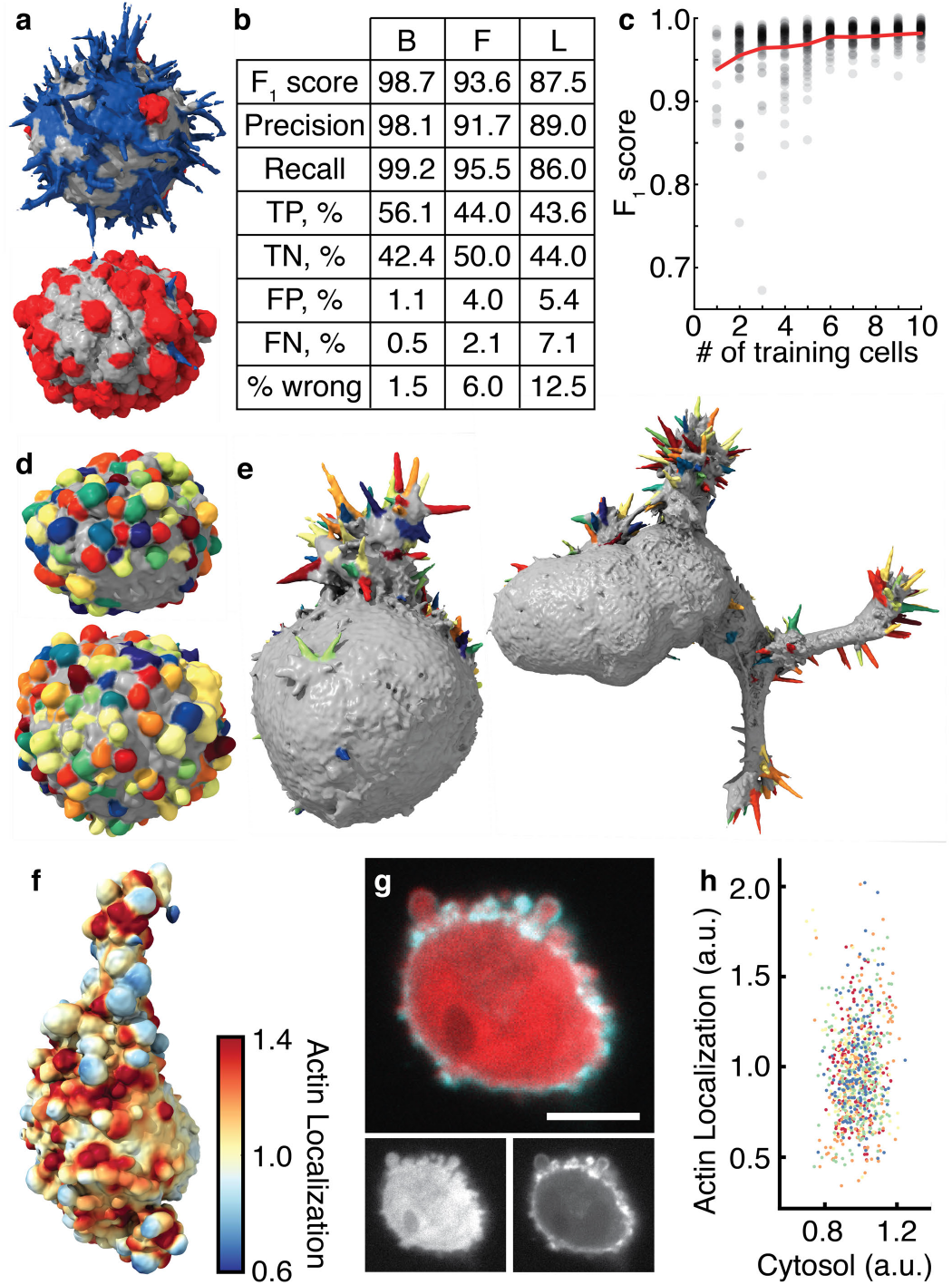
Validation and robustness of morphological motif detection. (**a**) A multiclass detector applied to cells derived from a human melanoma xenograft cultured in mice. Filopodia are shown blue, blebs are shown red, and areas with neither filopodia nor blebs are shown gray. (**b**) Validation measures for a bleb detector (left) trained on 19 MV3 melanoma cells, a filopodia detector (center) trained on 14 HBEC cells, and a lamellipodia detector (right) trained on 13 dendritic cells. TP, TN, FP, and FN are abbreviations for true positive, true negative, false positive, and false negative respectively. (**c**) The F_1_ score as a function of the number of cells trained on. The red line indicates the mean F_1_ score averaged over a maximum of 100 sets of cells, whereas the gray dots show individual sets of cells. (**d**) A bleb detector trained on the MV3 melanoma cell line applied to cells derived from a human melanoma xenograft cultured in mice. (**e**) A filopodia detector trained on xenograft-derived melanoma cells applied to HBEC cells. A filopodia detector trained on HBECs applied to the cell on the left is shown in Supplementary Fig. 5 and applied to the cell on the right is shown in Fig. 3. (**f**) An xy-slice of an MV3 cell expressing cytosolic GFP (red) and tractin-CyOFP (cyan). Scale bar, 10 μm. (**g**) Actin localization, measured over 1 μm, near the surface of an MV3 melanoma cell. Red indicates regions of relatively high localization, whereas blue indicates regions of relatively low localization. (**h**) Mean actin (tractin) localization vs. mean cytosolic localization on blebs. Each color indicates one of five different cells. The average localization of each cell in each channel was set to one.

### Validation and robustness of morphological motif detection

To validate the protrusion classification, we calculated the F_1_ score, which is the harmonic mean of precision and recall. Here, precision is defined as the ratio of patches correctly classified as protrusions to the total number of patches classified as protrusions, whereas recall is defined as the ratio of patches correctly classified as protrusions to the total number of patches that are protrusions. In calculating the F1 score, we only used patches selected by the trainer as certainly a protrusion or certainly not a protrusion. For four randomly chosen MV3 melanoma cells with extensive bleb formation, the F1 score calculated via leave-one-out-cross-validation over cells and averaged across three trainers was 0.986±0.006, corresponding to 1.3±0.6% incorrectly classified patches. This F1 score is likely high, in part, because only patches users were certain were blebs or non-blebs were included. We therefore also calculated F1 scores for the models where users clicked on all the blebs or all the non-blebs, yielding 0.77±0.03 and 0.76±0.04 respectively. However, as discussed above, these training data are biased toward selecting too few and too many blebs, respectively. Indeed, using these training data to validate our model, we find a 16±1% false positive rate (5±1% false negative rate) when users are asked to click on all the blebs and a 30±6% false negative rate (2±1% false positive rate) when users are asked to click on all the non-blebs. Validating over a larger number of cells with a single user, we measured an F1 score of 0.99 for 19 MV3 melanoma cells with blebs, 0.94 for 13 HBECs with filopodia, and 0.88 for 13 dendritic cells with lamellipodia (Fig. 4b).

An F1 score does not measure whether or not the workflow preferentially detects certain subtypes of protrusions. Since patch-merging algorithms could be sensitive to protrusion size, we used synthetic data to test the algorithm’s sensitivity to bleb size (Supplementary Fig. 6). On synthetic cells of radius 7.6 μm (48 pixels) we simulated blebs ranging in radius from 0.32 μm (2 pixels) and 0.64 μm (4 pixels) to 5.1 μm (32 pixels). Although only 70% of the smallest 0.32 μm radii blebs were decomposed as convex surface patches, almost all of the larger blebs were decomposed. A bleb detector trained on synthetic data correctly classified all blebs that were decomposed as convex surfaces.

Our workflow requires relatively little training data. We found that one user, training on just one cell in a dataset of 19 MV3 cells, yields an F1 score of 0.94±0.04 (mean ± standard deviation) on the remaining cells (Fig. 4c and Supplementary Fig. 7a). Increasing the training set size to 3 and 8 cells increases the F1 score to 0.97±0.01 and 0.987±0.004, respectively. Additional training data from one user improves the model accuracy relatively little, suggesting that models generated by a single user on different data sets would be similar. Indeed, models trained by a single user on distinct sets of four MV3 cells show 95.9±0.7% overlap, as measured by the Sorrenson-Dice index.^28^ This compares to an 88±3% overlap between models generated by different users (Supplementary Fig. 7b). These results underscore the importance of developing definitions of protrusive phenotypes that can be objectively and reproducibly applied to diverse datasets.

We found that models generated from training on one cell type can be extended to dissimilar cell types. Applying a bleb model generated from 19 MV3 cells, originating from a melanoma cell line, to 24 M405 cells, originating from a human melanoma xenograft cultured in mice, yields an F1 score of 0.97, which is the same score as the bleb model generated from the M405 cells applied to the M405 cells (Fig. 4d). The MV3 and M405 derived bleb models are also similar, showing 91% overlap when applied to the combined M405-MV3 dataset. To demonstrate that models can be successfully applied to entirely different cell types, we generated a filopodia model from 9 M405 melanoma cells, all but one of which exhibited only small numbers of filopodia. Applying this model to a dataset of 13 HBEC cells, which are transformed lung epithelial cells, yielded an F1 score of 0.90, compared to an F1 score of 0.96 applying this M405 model to the M405 cells (Fig. 4e). In conclusion, our morphological motif detector is nearly as accurate as manual data labeling, requires little training data, and, importantly, is portable between cell types, allowing objective comparisons between large numbers of diverse datasets.

### Association of morphological motifs with fluorescence signal distributions

To evaluate the ability of our morphological motif detector to be used to measure relationships between cell morphology, cytoskeletal organization, and intracellular signaling, we first examined the established relationship between actin localization and blebs.^29,30^ A bleb is initially devoid of actin, but as the bleb collapses it becomes enriched in actin. We sought to observe this well-known heterogeneity in actin localization on blebs. In a population of MV3 melanoma cells cultured in a 3D collagen matrix and expressing tractin-CyOFP^31^ as well as cytosolic GFP (Fig. 4f), we measured the localization of cytosol and actin (Fig. 4g) near the cell surface. To measure localization, at every mesh face we computed the average fluorescence intensity in a sphere around that face including only pixels within the cell. To correct for surface curvature-dependent artifacts, we depth-normalized the raw image prior to measuring localization.^32^ Comparing mean cytosolic intensity to mean actin intensity for each bleb, we found that, as expected, actin localization is more heterogeneous than localization of cytosolic GFP (Fig. 4h).

### Kras and PIP_2_ both associate with blebs, but do so differently

Equipped with a computational framework that permits the systematic and unbiased analysis of 3D cell morphology and molecular localization at the submicron scale, we set out to identify initial relationships between morphological motifs and signaling events. We again focused our analysis on blebs, since they are the predominant motif in melanoma cells in soft 3D environments^15^ and sought to measure how PIP_2_ and constitutively active Kras^V12^, may associate with blebs. Visually examining MIPs and individual slices of 3D images, both Kras^V12^ (Fig. 5a) and PIP_2_ (Fig 5b) appear to associate with blebs. To test this hypothesis, we measured the localization of Kras^V12^ within 2 μm of the cell surface for 13 MV3 melanoma cells (Fig. 5c). Applying a bleb detection model trained on MV3 cells expressing tractin-GFP, we found that cells expressing GFP-Kras^V12^ exhibited morphological structures, such as retraction fibers and uropods, which were not present in our actin-labeled dataset. The bleb detector sometimes classified these features as blebs (Supplementary Fig. 1a), which could lead to bias. To exclude those structures from the analysis we built a retraction fiber/uropod detector (Supplementary Fig. 8b,c) and subtracted those patches from the set of detected blebs (Supplementary Fig. 8d). Measuring the mean Kras^V12^ localization on and off detected blebs, we found no statistically significant difference (p-value: 0.5, effect size: -0.006, t-statistic: -0.017, Fig. 5d,e). However, measuring Kras^V12^ localization as a function of distance from a bleb edge, we found that Kras^V12^ does localize to bleb edges (Fig. 5f). In contrast, a population of 35 cells expressing GFP in the cytosol and with intensity also measured within 2 μm of the cell surface, showed no localization to bleb edges (Fig. 5f). Similarly, measuring bleb density locally over a scale less than that of a single bleb, we observed that high Kras associates with intermediate local bleb densities, which are near bleb edges, whereas low Kras associates with both low and high local bleb densities, which are not near bleb edges (Supplementary Fig. 9a-c). Simulating uniformly labeled surface distributions in synthetic cells, we demonstrated that intensity localizes to bleb edges for a variety of surface thicknesses, but not for cytosolically labeled cells (Supplementary Fig. 10a). Kras^V12^ intensity localization at bleb edges is therefore consistent with a uniform cortical signal distribution.

**Figure 5.**
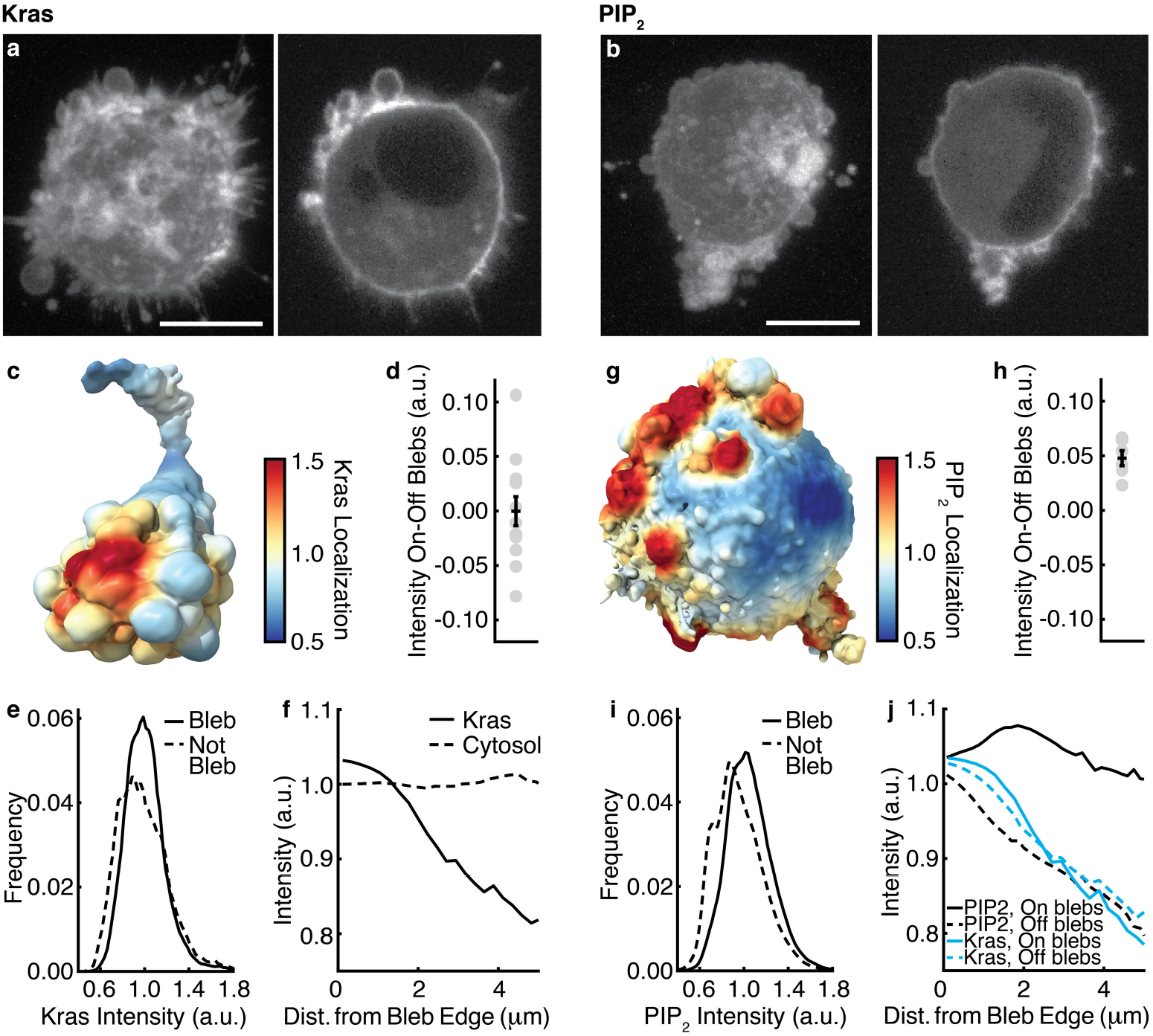
Kras and PIP_2_ associate with blebs differently. (**a**) An MV3 expressing GFP-Kras^V12^ cell shown as a MIP (left) and an xy-slice (right). (**b**) An MV3 cell expressing PLCΔ-PH-GFP shown as a MIP (left) and an xy-slice (right). (**c**) Kras localization, measured over 2 μm, near the surface of an MV3 cell expressing GFP-Kras^V12^. (**d**) The differences between the mean Kras intensity on and off blebs for 13 cells. (**e**) Distributions of Kras intensity for mesh faces on and off blebs for the same cells as in **d**. (**f**) Fluorescence localization vs. distance from a bleb edge for 13 GFP-Kras^V12^ labeled cells and 35 GFP cytosolically labeled cells. (**g**) PIP_2_ localization, measured over 2 μm, near the surface of an MV3 cells expressing PLCΔ-PH-GFP. (**h**) The differences between the mean PIP_2_ intensity on and off blebs for 6 movies of cells. (**i**) Distributions of PIP_2_ intensity for mesh faces on and off blebs for the same cells as in **h**. (**j**) PIP_2_ and Kras localization, both on and off blebs, vs. distance from a bleb edge. Scale bars, 10 μm.

Like Kras^V12^, PIP_2_ localizes to the plasma membrane, however, unlike Kras^V12^, PIP_2_ associates with blebs. Measuring the localization of PIP_2_ within 2 μm of the cell surface for 6 movies of MV3 cells expressing PLCΔ-PH-GFP (Fig 5g), a PIP_2_ translocation biosensor, we found that PIP_2_ localizes to blebs (p-value: 0.0005, effect size: 1.7, t-statistic: 6.9), with each cell having a higher mean intensity on blebs than off (Fig. 5h and 5i). Interestingly, cytosolically labeled GFP showed less localization to blebs than non-blebs both by visual inspection and by quantitative measurement (Supplementary Fig. 10b), possibly due to slow diffusion of the free probe into blebs. Consistent with a surface fluorescence distribution, PIP_2_ like Kras^V12^ also associates with bleb edges (Fig. 5j). However, whereas Kras^V12^ localization falls off with increasing distance from a bleb edge both on and off blebs, PIP_2_ localization falls off with increasing distance from a bleb edge only off blebs. Similarly, measuring the local bleb density we found that high PIP_2_ associates with intermediate local bleb densities, which are near bleb edges (Supplementary Fig. 9d). Low PIP_2_ associates with small local bleb densities, which are away from blebs, but not with large local bleb densities, which are on blebs. In conclusion, visual inspection of 3D images suggests that Kras^V12^ and PIP_2_ similarly associate with blebs (Fig. 5a,b). However quantitative analysis shows that their association with blebs obeys fundamentally different mechanisms.

Our workflow supports many other types of analyses relating cell morphology and molecular distributions. For example, underlying the extraction of morphological motifs is a rich quantification of geometric properties. Based on 3D surface renderings of PIP_2_ localization (Fig. 6a), we hypothesized that PIP_2_ may preferentially associate with large blebs. Measuring bleb volume, we found that indeed larger blebs show greater association with PIP_2_ than smaller blebs (Fig. 6b). This contrasts to cytosolically labeled cells, where no such relationship is found (Supplementary Fig. 10c). Our workflow also measures the cellular-scale configuration of morphological motifs using spherical statistics. Analyzing the average direction of blebs and PIP_2_ localization relative to the cell center, we found that blebs and PIP_2_ co-polarize (Fig. 6c). Since the cell is not a sphere, as assumed in spherical statistics, we constructed control distributions by randomizing the population of patches classified as blebs. Since the study of many signaling pathways benefit from measuring not just morphology, but also morphodynamics, we also developed a measure of boundary motion at each mesh face. Fig. 6d shows the boundary motion of a PIP_2_-labled cell over several frames. Measuring the motion difference over ~30 sec, which is on the order of the bleb lifetime,^33^ we found that blebs preferentially associate with regions of protrusive motion (Fig. 6e,f). We also observed that regions of high PIP_2_ tend to be more retractive than regions of low PIP_2_ (Fig. 6g), which is consistent with increased PIP_2_ localization on blebs because blebs form and retract cyclically.

**Figure 6.**
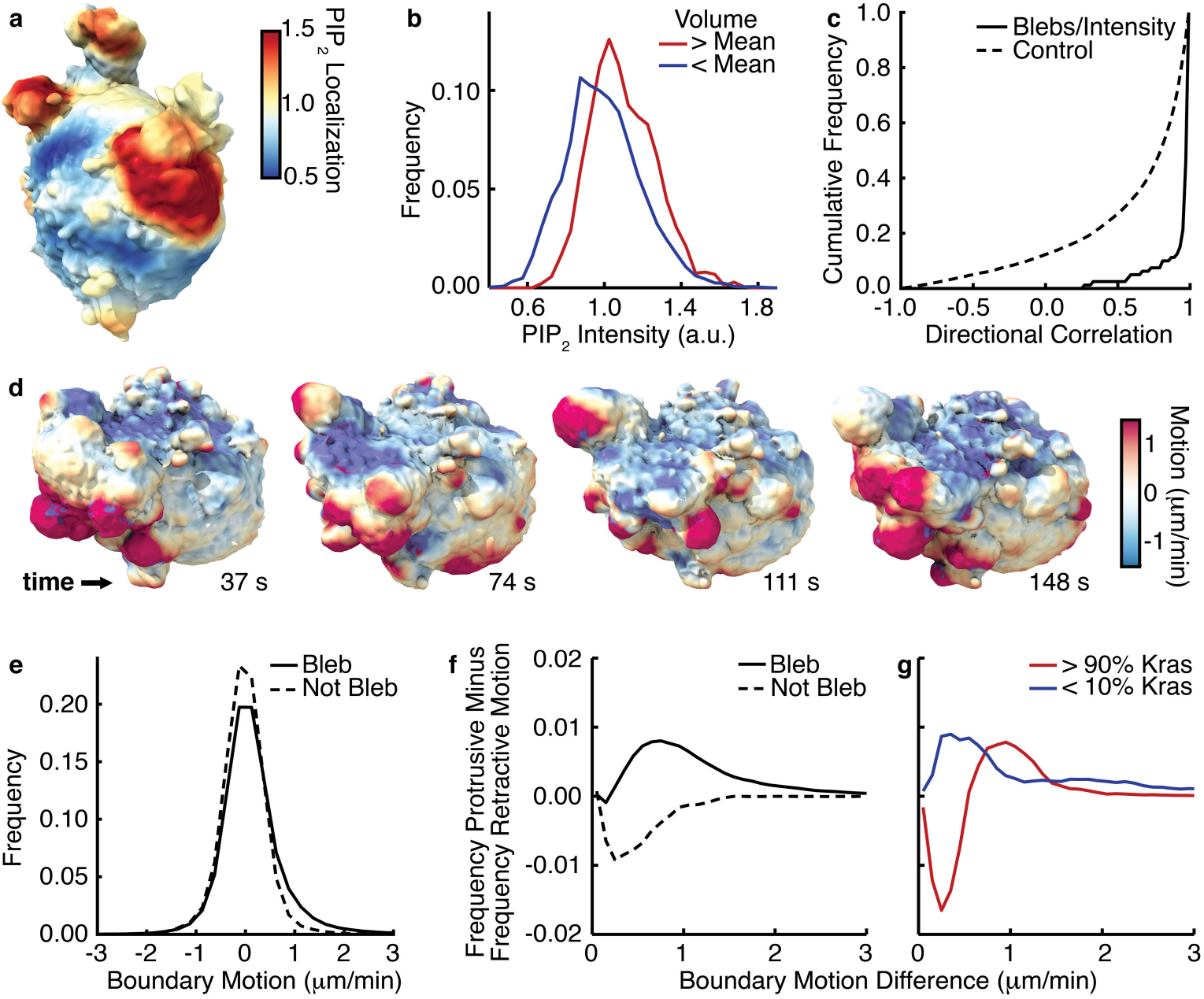
Further PIP_2_ analyses. (**a**) PIP_2_ localization, measured over 2 μm, near the surface of an MV3 cell expressing PLCΔ-PH-GFP. (**b**) Distributions of PIP_2_ intensity for mesh faces on blebs of greater than average volume and on blebs of less than average volume. (**c**) The directional correlation of blebs with PIP_2_ localization. The cumulative correlation distribution is shown in purple, and the cumulative control distribution is shown in gray. (**d**) Surface renderings of the boundary motion of an MV3 cell. Red indicates regions of high protrusive motion, whereas blue indicates regions of high retractive motion. (**e**) Boundary motion distributions at mesh faces on and off blebs. (**f**) The frequency of protrusive motion minus the frequency of retractive motion on and off blebs as a function of surface speed, (**g**) The same measure shown in **f** for mesh faces in the top and bottom deciles of PIP_2_ localization.

## Discussion

Recent advances in high-resolution 3D light-sheet microscopy,^11-16^ allow the direct observation of subcellular processes at unprecedented spatial and temporal scales. However, incorporating these observations into a scientific framework that enables data exploration, hypothesis testing, and ultimately the development of new biological theories is a challenge that we have only begun to undertake. Even simply visualizing a 3D image on a 2D screen requires imposing an interpretation on it.^19^ Compared to 1D and 2D features, such as length and area, human observers also exhibit decreased ability to make quantitative comparisons of 3D features, such as volume.^34^ Additionally, the scale of data presents hurdles to interpretation. A single movie from a light-sheet microscope can be thousands of frames long and 1 TB in size.^19^ Such large data sizes not only prohibit intensive examination of individual frames, but also tax even state-of-the-art computing infrastructures. To interpret light-sheet microscopic data, further development of 3D computing infrastructures and image analysis algorithms is required.

Here, we focused on developing algorithms that enable the analysis of biological surfaces. In particular, we developed an algorithm to detect diverse morphological motifs on the 3D cell surface using a machine learning approach. This offers the flexibility to quantify the frequency, configuration, and geometry of any motif of interest. As a demonstration, we trained classifiers to detect blebs, filopodia, and lamellipodia, which are three abundant morphological motifs found on cells. To detect a new type of morphological motif, users need only click on examples of surface regions that are and are not that motif. Our algorithm speeds up analysis by relieving the user from attempting to count motif occurrences in 3D data, which can be extremely difficult and is often effectively impossible. It also enables objective analysis, not only via complete and unbiased measurement of morphological parameters within a single dataset, but also between perturbations, projects, and laboratories. This is the first generic morphological motif detector and is one of the first image analysis tools for cell biology that incorporates techniques from computer graphics. Clearly, with the rapid rise of 3D microscopy in live cell and tissue imaging, computer graphics methods will become an important factor in biological discovery.

In addition to a morphological motif detector, we developed and integrated a suite of tools for investigating the coupling between morphology, morphodynamics and molecular-scale mechanisms. As a proof of concept, we used these tools to examine the differential association of Kras^V12^ and PIP_2_ signaling with cell surface blebs. We expect that our tools will become the basis for projects ranging from cell behavioral screens and cell migration studies, to molecularly specific investigations of signal distribution. Although we validated our methods for single cells imaged via advanced light-sheet microscopy with nearly isotropic resolution,^11-16^ many of our analyses could likely be extended to other imaging modalities following independent validation. In particular, our algorithms could in principle be applied to the analysis of surfaces of multicellular biological structures, such as spheroids and organs.

## Acknowledgements

This research was funded by grants from the Cancer Prevention Research Institute of Texas (RR160057 to RF and R1225 to GD) and the National Institutes of Health (F32GM116370 and K99GM123221 to MKD, K25CA204526 to ESW, F32GM117793 to KMD, and R01GM067230 to GD). Most surface renderings were performed in UCSF ChimeraX, which was developed by the Resource for Biocomputing, Visualization, and Informatics at the University of California, San Francisco (supported by P41GM103311). We thank Tom Goddard for assistance with ChimeraX, as well as Ingrid de Vries, Jörg Renkawitz, and Michael Sixt for assistance differentiating dendritic cells. We would also like to thank Peter Friedl for the MV3 melanoma cells, Sean Morrison for the primary melanoma cells, John Minna and Jerry Shay for the transformed HBEC cells, and Michael Sixt for the dendritic cell precursors.

## Methods

### Cell culture and labeling

All cells were cultured at 5% CO2 and 21% O2. MV3 melanoma cells (a gift from Peter Friedl at MD Anderson Cancer Center) were cultured using DMEM (Gibco) supplemented with 10% fetal bovine serum. Primary melanoma cells (a gift from Sean Morrison at UT Southwestern Medical Center) were cultured using the Primary Melanocyte Growth Kit (ATCC). Human bronchial epithelial cells (HBEC; a gift from John Minna at UT Southwestern Medical Center), immortalized with Cdk4 and hTERT expression and transformed with p53 knockdown, Kras^V12^, and cMyc expression,^35^ were cultured in keratinocyte serum-free medium (Gibco) supplemented with 50 mg/ml of bovine pituitary extract (Gibco), 5 ng/ml of EGF (Gibco), and 1% Anti-Anti (Gibco). Conditionally immortalized hematopoietic precursors to dendritic cells^36^ that express Lifeact-GFP^37^ (a gift from Michael Sixt, IST Austria) were cultured and differentiated as previously described.^38^

Fluorescent constructs were introduced into cells using the pLVX lentiviral system (Clontech) and selected using antibiotic resistance to either puromycin or geniticin. The GFP-tractin construct contains residues 9–52 of the enzyme IPTKA^39^ fused to GFP.^40^ The CyOFP-tractin peptide contains the tractin peptide fused to the CyOFP protein. CyOFP is a cyan-excitable orange fluorescent protein with peak excitation at 505 nm and peak emission at 588 nm.^31^ The GFP-Kras^V12^ plasmid was constructed by cloning a Kras^V12^ fragment from the pLenti-Kras^V12^ construct^35^ into the pLVX-GFP vector. The biosensor for PIP_2_, PLCΔ-PH-GFP, encodes a PI(4,5)P2 lipid selective PH domain that can be used as a fluorescent translocation biosensor to monitor changes in the concentration of plasma membrane PI(4,5)P2 lipids.^41^ Some MV3 cells expressing GFP in the cytosol, which were analyzed here as a control population, appeared in a previous publication.^15^

Collagen gels were created by mixing bovine collagen I (Advanced Biomatrix) with concentrated PBS and water to a collagen density of 2.0 mg/ml. This collagen solution was then neutralized with 1N NaOH and mixed with cells just prior to incubation at 37 °C to induce collagen polymerization.

### Light-sheet imaging

Imaging was performed via microenvironmental selective plane illumination microscopy,^15^ a type of two-photon Bessel beam light sheet microscopy that confers near-isotropic resolution (300 nm lateral, 340 nm axial) and permits recording of cell behavior several millimeters from mechanically perturbing hard surfaces. Images were acquired at 37 °C in a non-descanned image capture mode with an axial step size of 160 or 200 nm and an excitation wavelength of 900 nm. Melanoma cells were imaged in cell culture medium supplemented with HEPES buffer to maintain pH during imaging.

### Image deconvolution

As a first step towards analysis, we Wiener deconvolved each 3D image as previously described.^15^ Unless otherwise specified, all microscopy images shown here are raw, non-deconvolved images. The Wiener parameter, which is the inverse of the signal-to-noise ratio, was usually set to 0.018. To better detect the dim ends of filopodia, for filopodia detection it was set to 0.015. For cytosolically labeled cells, we automatically estimated the parameter in each frame by defining the signal as the average fluorescence intensity within the cell and the noise as the standard deviation of the fluorescence intensity outside the cell. Following deconvolution, an apodization filter was applied to the optical transfer function (OTF) of the image in the spatial frequency domain. This filter had a value of 1 at the origin and decayed linearly to 0 at the edge of the filter support, which is set by the user as a percentage of the maximum OTF value. This threshold value, here termed the apodization height, was usually adjusted according to the homogeneity of the fluorescence label and the fineness of the morphological motif being detected. Higher apodization heights smooth the image more and allow for more robust detection of large objects, whereas lower apodization heights allow for the detection of finer structures but also admit more noise. For lamellipodia and bleb detection on Kras and PIP_2_ labeled cells, it was set to 0.06, for bleb detection on tractin and cytosolically labeled cells, it was set to 0.04, and for filopodia detection it was set to 0.

### Cell surface extraction

The deconvolved images were further processed prior to cell surface extraction. For bleb detection on tractin and cytosolically labeled cells, an Otsu threshold was first calculated from the 3D image,^42^ holes were then filled using a 3D grayscale flood-fill operation, and objects disconnected from the main cell were removed. Matlab’s *isosurface* function was finally used to create a triangle mesh at the intensity value specified by the Otsu threshold.

For filopodia detection, we similarly extracted the cell surface, however the reduced apodization compared to bleb detection yielded a noisier image, which we compensated for by smoothing the deconvolved image with a 3D Gaussian kernel of standard deviation 0.6 pixels.

For bleb detection on Kras and PIP_2_ labeled cells, we processed the deconvolved images similarly to tractin and cytosolically labeled cells used for bleb detection. However, since Kras and PIP_2_ label the cell less homogeneously, we first applied a gamma correction of 0.7 to the deconvolved images. Even with this correction, in PIP_2_ labeled cells, the nucleus was sometimes not segmented along with the cell. For all PIP_2_ labeled cells, we therefore also combined the gamma corrected image with an “inside” image that segmented the cell interior. To create the “inside” image from the gamma corrected image, we applied an additional gamma correction of 0.6, smoothed the image with a 3D Gaussian kernel of width 2 pixels, Otsu thresholded the image, morphologically dilated the image by 4 pixels, filled holes in each xy-slice, morphologically eroded the image by 6 pixels, and finally smoothed the binary image with a 3D Gaussian kernel of width 1 pixel. Since this process shrinks the cell, if the parameters are chosen correctly the edges of the morphological motifs should mostly lay outside the “inside” image. To combine the “inside” image with the deconvolved image, we normalized the deconvolved image by its Otsu threshold value, took the pixel-by-pixel maximum of this image and the “inside” image, and extracted a triangle mesh as an isosurface at an intensity level of 1.

The ends of the long, thin lamellipodia of dendritic cells fail to segment using the techniques described above. To better segment lamellipodia, we combined the “inside” and normalized deconvolved images described above for PIP_2_ labeled cells with a “surface filtered” image that enhances planar features, such as lamellipodia (Supplementary Fig. 2). The surface filter, which was developed by Elliott *et al*.,^32^ uses multiscale Gaussian second order partial-derivative kernels of the form

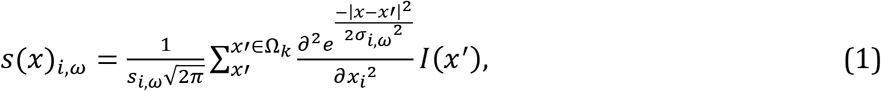

where *I*(*x*) is the image intensity, *σ_i,ω_* is the half width of the Gaussian in dimension *i* at scale *ω*, Ω_*k*_ is the filter kernel support, and *s*(*x*)_*ω*_ is the filter response at scale *ω*. The total filter response, *S*(*x*), is merged across scales via

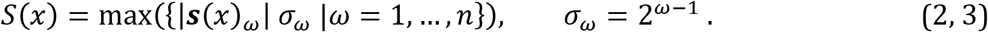

We used filter scales 1, 1.5, 2, and 4 pixels to segment lamellipodia of various thicknesses. To combine the response of the surface filter with the “inside” and normalized deconvolved images, we normalized the response by subtracting both the mean image intensity and twice the standard deviation of the image intensity prior to dividing by the standard deviation of the image intensity. The resultant triangle meshes were smoothed using 6 iterations of curvature flow smoothing.^43^

Although not used in this paper, our software also includes the option to segment cells by combining a normalized deconvolved image with a steerable filtered image. Steerable filters are computationally efficient edge detectors that, depending upon the parameters chosen, enhance linear or planar structures at specified scales.^44,45^

Segmentations were spot checked by thresholding the 3D image at the isosurface intensity value immediately prior to mesh extraction and examining the overlaid raw and thresholded images as 3D image stacks in ImageJ^46^ (Supplementary Fig 11). For analyses where internal mesh cavities could alter results, meshes were also exported to ChimeraX^47^ for further examination. Segmentations that were found to be inaccurate or had cavities were excluded from further analysis.

### Decomposition of the cell surface into convex patches

Although the image deconvolution and cell surface extraction parameters require customization for different cell types, the remainder of the workflow does not, and its parameters were kept constant throughout the paper.

To decompose the cell surface into convex patches, we first performed a watershed segmentation of surface mean curvature, as previously described.^15^ This oversegments the cell surface into small patches, which are analogous to superpixels in image analysis, which we later merge to create convex patches. First, we calculated the mean and Gaussian curvature at every triangle face.^32,48^ Next, we constructed an adjacency graph of faces where each face is a node that is connected to exactly three other spatially adjacent faces. Matlab’s *isosurface* function does not always produce triangle meshes with sufficient topological consistency to create such a graph. Our software fixes common topological inconsistencies, such as triangular edges that are only connected to one face. Rarely, however, a face graph cannot be constructed. In these situations, very slightly changing the image deconvolution parameters usually solves the problem, although we did not need to do so here. Since curvature can be noisy, we next smoothed mean curvature in two different ways. First, we used a kd-tree to median filter curvature in 3D space over 2 pixels. The meSPIM is Nyquist sampled, and so 2 pixels, which is 320 nm, is approximately the microscope’s spatial resolution. Second, to reduce spurious curvature fluctuations, we diffused mean curvature on the mesh using a diffusion kernel^49,50^ according to the equation

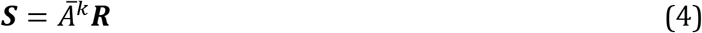

for 20 iterations, where ***R*** is the curvature, 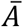 is a normalized, weighted adjacency matrix of the faces graph, *k* is the number of iterations, and ***S*** is the smoothed curvature. We defined *A* as

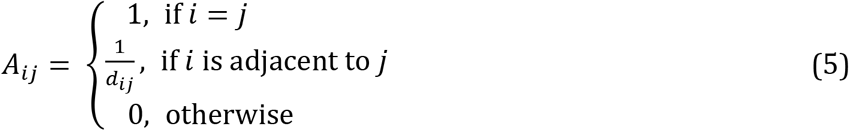

where *d_ij_* is the distance between faces *i* and *j*. To normalize *A*, we multiplied it by a diagonal matrix, where each diagonal element was the inverse of the sum of that row. Next, we performed a watershed segmentation of the smoothed curvature over the cell surface.^24^ Watershed segmentations are often performed on 2D images, where each pixel is adjacent to exactly four other pixels. Here, we similarly performed a watershed segmentation over the adjacency graph of faces, where each face is adjacent to exactly three faces.

We next merged adjacent patches using a spill depth criterion.^24^ Here, the spill depth between two adjacent patches was defined as the maximum curvature of the two patches minus the maximum curvature at the patch-patch interface. This is analogous to the depth of water that the patch can hold before spilling into the neighboring patch. Starting with the smallest spill depth, we merged patches until no spill depth was below a cutoff of 0.6 times the Otsu threshold of mean curvature for the cell. Supplementary Fig. 3 shows the effect of altering the spill-depth cutoff and other patch merging parameters.

Finally, we decomposed the surface into approximately convex patches by iteratively applying the triangle and line-of-sight (LOS) criteria. To apply the triangle criterion,^15^ we first calculated the closure surface area for each patch and pair of adjacent patches. We defined the closure surface area as the minimum additional surface area needed to create a closed polyhedron from a surface patch. We then merged adjacent patches if they meet the criterion

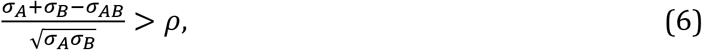

where *σ_A_* and *σ_B_* are the closure surface areas of the two patches, *σ_AB_* is the closure surface area of the merged patch, and *ρ* is the triangle cutoff parameter, which we here set to 0.7. The triangle criterion can be thought of as similar to the law of cosines and intuitively seeks to merge patches that meet at small angles. Starting with the largest *ρ*, we merged all pairs of patches that met the triangle criterion before applying the LOS criterion.

The LOS criterion merges adjacent patches with high mutual visibility.^25,51^ We defined the mutual visibility of patches *A* and *B* as the percentage of line segments that connect a face in *A* with a face in *B* that are lines of sight, where a line of sight is a line segment that falls entirely within the mesh. We calculated mutual visibility by randomly selecting a face on each patch, and using a triangle-ray intersection^52^ algorithm to determine whether a line segment connecting the two faces exited and reentered the mesh. A small patch and an adjacent very large patch may have a large mutual visibility because of lines of sight that extend across the width of the cell, even if these two patches should not be merged. When merging two patches, we therefore discarded line segments that were longer than twice the smaller patch size. Supplementary Fig. 12a shows the convergence of mutual visibility as a function of the number of line segments tested. We calculated mutual visibility from 20 line segments per pair of patches. In an exact convex decomposition, any two points within any patch could be connected by a line of sight. However, because of biological variation and image noise, requiring a mutual visibility of 1 is too strict a requirement for cell images. We instead merge patches if their mutual visibility is greater than 0.7. Starting with the largest mutual visibility between patch pairs, we merged all patch pairs meeting the LOS criterion, before again applying the triangle criterion.

Having three patch merging criteria for convex surface decomposition allows us to balance accuracy, speed, and robustness to noise. The spill-depth criterion is fast but potentially inaccurate, whereas the LOS criterion is relatively slow, but exact. The triangle criterion implements the short-cut rule,^26^ which biases merging towards certain types of convex decompositions. By adjusting the three merging parameters, users can control which criteria dominate in their analysis.

### Classification of morphological motifs

To classify each patch by morphological motif, we first performed feature selection on the geometric patch features listed in Table 1. Implemented by the Matlab built-in function *sequentialfs()*, our sequential feature selection randomly successively removed features as long as doing so reduced the misclassification rate. The misclassification rate was measured using 10-fold cross validation. The geometric features selected can vary considerably from dataset to dataset even for similar training sets, presumably because of correlations between features, randomness, and dataset differences. For example, Supplementary Table 4 shows the features selected for bleb detection models generated by three different users training on the same four cells. In this example, no feature was selected by all three models, and no two models shared more than two selected features. Once features were selected, features were normalized to have the same mean and standard deviation, and a linear support vector machine (SVM) was used to classify patches. Since SVM models vary from user to user, to analyze actin, Kras, and PIP_2_ localization, we had models created by three different users vote on the classification of each bleb.

### Characterization of patches

To classify patches by morphological motif, we calculated geometric descriptions of each patch. The full list of 23 features used by the SVM classifier is provided in Table 1. In calculating these features, mean curvature was smoothed as described above, but Gaussian curvature was not. We defined the average patch position as the mean location of the faces in the patch, and we similarly defined the weighted average patch position as the mean location of the faces weighted by curvature. The feature ‘variation from a sphere’ was defined by the standard deviation of the distances from a patch s faces to the average patch position divided by the mean distance of those faces to the average patch position. We defined the closure surface area as described above. The closure center was also defined as the mean position of the mesh vertices at the patch edge. We defined the patch radius as the mean distance of the patch s faces from the closure center.

The volume, *V*, was calculated using the equation

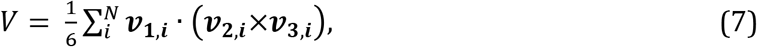

where *N* is the number of faces, and ***ν***_***1,i***_,***ν***_***2,i***_ and ***ν***_***3,i***_ are the vertices of face *i*. The vertices must be ordered such that the face normal extends outwards from the cell. To derive this equation, the mesh can be thought of as decomposed into tetrahedrons where the vertices of each tetrahedron are those of a face combined with the origin.^53^ The signed volumes of the tetrahedrons sum to the volume of the mesh. Patches were closed prior to calculating their volumes.

We calculated the shape diameter function similarly to Shapira *et al*.^27^ For each patch, we randomly picked 20 mesh faces on the patch and extended a ray inwards from the mesh face at a randomly chosen angle within *π*/3 of the direction opposite to the face’s normal. We calculated the distance each ray traveled before intersecting the opposite side of the mesh. The shape diameter function of the patch was then defined as the mean travel distance within one standard deviation of the median distance.

### Optional merging of convex patches

Some morphological motifs, such as lamellipodia and flagella, are not convex but are composed of multiple convex regions. To detect such motifs, we optionally merge convex patches into patch composites. Since adjacent patches that compose a larger structure often have smooth curvature at their interface, we first merge patches using a modified line of sight criterion with line segment length capped at 10 pixels and a mutual visibility cutoff of 0.7. The line of sight criterion is described above. This step is not required for convex patch merging and can be disabled by the user. We next employed a more versatile machine learning based framework to merge adjacent patches. Using the geometric features for pairs of adjacent patches listed in Table 2, as well as user provided training data, we trained an SVM to automatically merge patches. We used the same feature selection procedure and SVM parameters as for patch classification.

### Characterization of adjacent patches

To merge adjacent patches into patch composites using an SVM, we calculated geometric characterizations of each pair of adjacent patches. The full list of 36 features used by the SVM is provided in Table 2. Some measures of patch pairs incorporate individual patch measures, which are described above. Unless otherwise specified, mean curvature was smoothed as described above, but Gaussian curvature was not.

To better describe the surface geometry at patch-patch interfaces, we calculated the two principal curvatures, *κ*_1_ and *κ*_2_, everywhere on the cell surface,

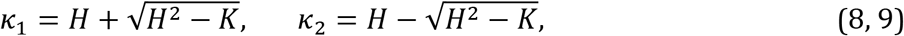

where *H* is the unsmoothed mean curvature and *K* is the unsmoothed Gaussian curvature. For various geometries defined by principal curvature values, we then calculated the fraction of the interface that had that geometry. As a noise threshold, we used the standard deviation of the smoothed mean curvature. Principal curvatures above this threshold or below the negative of this threshold were defined as large, and those more than four times above or below it as very large. We defined a ridged geometry as a large positive *κ*_1_and a small *κ*_2_, a very ridged geometry as a very large positive *κ*_1_ and a small *κ*_2_, a valley-like geometry as a small *κ*_1_ and a large negative *κ*_2_, a very valley-like geometry as a small *κ*_1_ and a very large negative *κ*_2_, a domed geometry as a large positive *κ*_1_ and a large positive *κ*_2_, a cratered geometry as a large negative *κ*_1_ and a large negative *κ*_2_, a flat geometry as a small *κ*_1_ and a small *κ*_2_, and a saddle-like geometry as a large positive *κ*_1_ and a large negative *κ*_2_.

### Generation of training data

We designed a graphical user interface to enable the collection of training data necessary for motif classification. Users are shown a surface rendering of a cell with surface patches outlined and can interact with the cell by rotating and moving it, and zooming in and out on regions of interest. To generate data for patch classification, we asked users to click on patches that are certainly the morphological motif of interest and then subsequently asked them to click on patches that are certainly not that motif. Similarly, to generate data for the optional step of convex patch merging, we asked users to click on pairs of patches that should certainly be merged and then asked them to click on pairs of patches that should certainly not be merged. Pairs of patches that were not adjacent were automatically excluded from the training set. We have successfully tested this interface in Matlab versions R2017b and R2013b. However, since in Matlab user interface functionality can vary from version to version, it may not work in some versions of Matlab.

### Generation and analysis of synthetic images

For algorithm validation, we created synthetic spherical cells of radius 48 pixels. The cell size was chosen to mimic the pixel spacing on the meSPIM of 0.16 μm per pixel for a cell 7.6 μm in radius. Placed randomly on the cells’ surfaces were spherical blebs that ranged in radius from 2 to 32 pixels and in number from 4 to 256 per cell. (See Supplementary Fig. 6 for example synthetic cells). Since pixelation at the cell edge could hamper the cell surface extraction and subsequent analysis, edge pixels were subdivided into a finer 3D grid to calculate the percentage of the pixel occupied by the synthetic cell. The final synthetic images were saved with 32 grayscale intensity values. Synthetic cells were not deconvolved, but the remainder of the analysis workflow was identical to that used for microscopic data. The same surface extraction parameters were used as for bleb detection on tractin and cytosolically labeled cells.

### Mapping fluorescence intensity to the cell surface

To measure the fluorescence intensity local to each mesh face, we used the raw, non-deconvolved, fluorescence image. At each mesh face, we used a kd-tree to measure the average pixel intensity within the cell and within a sampling radius of the mesh face. To correct for surface curvature dependent artifacts, we depth normalized^32^ the image prior to measuring intensity localization by normalizing each pixel by the average pixel intensity at that distance interior to the cell surface. Prior to analysis, we also normalized each cell’s surface intensity localization to a mean of one.

### Calculation of distance from a bleb edge

On the adjacency graph of faces, we calculated the distance from each face to the nearest bleb edge measured in number of faces traversed. To convert this distance to micrometers, we multiplied by the average distance between faces for each cell in each frame. Since the distance in micrometers between adjacent faces varies, our calculation of distance is an estimate rather than exact.

### Calculation of local bleb density

To calculate bleb density, we first assigned the value one to each mesh face on a bleb and the value zero to each mesh face not on a bleb (Supplementary Fig. 9a). We then diffused these values on the mesh surface using Eq. 4 over 600 iterations (Supplementary Fig. 9b). We choose the number of iterations such that the bleb density would be calculated over a short distance on the order of a bleb length. Eq. 4 does not allow an exact measurement of bleb density and may be unstable over distances on the order of many bleb lengths.

### Spherical statistics

The von Mises-Fisher distribution is defined on an 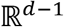 sphere within 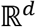 space.^54^ For *d* = 2 dimensions it approximates a wrapped normal distribution on a circle, and, similar to the normal distribution, for any *d* is parameterized by a mean and an inverse spread. For *d* = 3 dimensions, the von-Mises Fisher distribution is

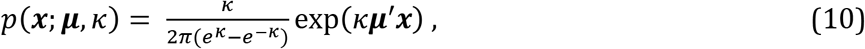

where ***μ*** is the mean direction parameter and *κ* is the concentration parameter, which is inversely related to the data spread. The maximum likelihood estimate of the mean direction is simply

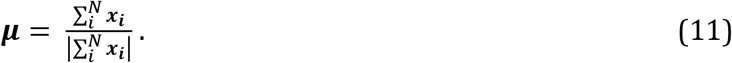

A Newton’s Method approximation for *κ, κ*_2_, in three dimensions is

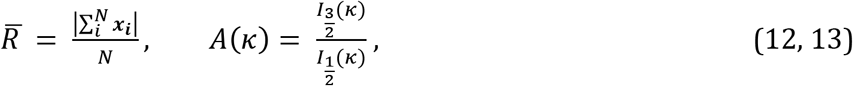

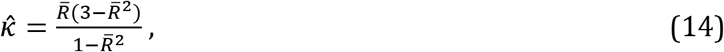

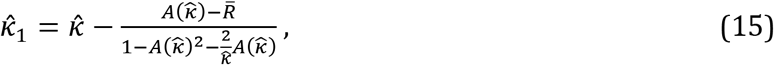

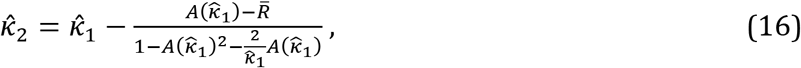

where *N* is the number of data vectors and *I* are Bessel functions of the first kind.^54^

In Fig. 6f we computed the directional correlation of morphological motifs, here blebs, with intensity localization. In each frame, we defined the directional correlation as ***μ_blebs_ · μ_intensity_***. To measure ***μ_blebs_***, we calculated a set of unit vectors, ***x_blebs_***, that extended in the direction from the cell center, defined as the location of the pixel farthest from the cell edge, to each mesh face on a bleb. To measure ***μ_intensity_***, we calculated a set of unit vectors, ***x_intensity_***, that extended in the direction from the cell center to every mesh face, and in Eq. 11 we weighted ***x_i_*** by the intensity localization. Since the cell is not a sphere and most cells have polarized shapes, the surface itself is expected to have a nonrandom ***μ*** and a small *κ*. To account for this, we created a control distribution of directional correlations ***μ_blebsRand_ · μ_intensity_***, where ***μ_blebsRand_*** was calculated from a set of vectors where the patch classification was randomly permuted. In each frame, we created 200 such permutations by randomly assigning patches to be a bleb or not a bleb. Although our study does not include analysis of *κ*, the software package computes *κ* for future analyses and thus is documented here for the sake of completeness.

### Measurement of boundary motion

To measure boundary motion, for each face we found the closest face in the previous frame using a kd-tree. We then defined the boundary motion as

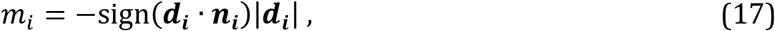

where *m_i_* is the boundary motion at face *i*, ***d_i_*** is the vector from face *i* to the closest point in the previous frame, and ***n_i_*** is the normal to the surface at face *i*.

This is not an ideal measure of boundary motion since the mapping vectors ***d_i_*** may cluster on select faces of the previous frame’s surface, or even alter the topology among faces, in a physically unrealistic manner (see Machacek *et al*.^55^ for an illustration of these problems with 2D boundaries). As a control, we also calculated the boundary motion for each face by finding the closest point in the next frame. Supplementary Fig. 12b shows the protrusive and retractive motion of six cells using both definitions of boundary motion. Here, backwards motion is the mapping of points from each frame to the previous frame and is the definition used elsewhere, and forwards motion is the mapping of points from each frame to the subsequent frame. Even though the backwards and forwards motions of the cells are different, in both cases blebs are more protrusive than non-blebs. This measure is also not a subpixel measure of motion, and should not be used to measure subpixel motions. Because we map each face to the closest face rather than the closest surface point in the previous frame, motions that are less than the average distance between faces will be undersampled in the motion distribution.

### Statistical hypothesis testing

For each Kras and PIP_2_ labeled cell, we measured the mean intensity localization of faces on and off blebs and then performed a one-sided t-test on the differences of the means after testing for normality using a Kolmogorov-Smirnov test. The Cohen’s d effect size was measured.

Unless otherwise indicated, all errors and error bars show the standard error of the mean.

### Surface rendering

The majority of triangle meshes were rendered in ChimeraX.^47^ Colored triangle meshes were exported from Matlab as Collada.dae files using custom-written code and were rendered using full lighting mode. Lighting intensity and ambient intensity were adjusted. Colormaps were modified from colorBrewer.^56^ The surfaces in Supplementary Figures 5, 7, and 10 were rendered within Matlab. Our software is capable of rendering all meshes shown in the paper within Matlab, as well as creating Collada files for export to ChimeraX.

### Data availability

Some of the data analyzed here is provided with the software for demonstration purposes. The remainder of the data that support the findings of this study are available from the corresponding author upon reasonable request.

### Software availability

The software described here, as well as a user’s guide and test data will be made available upon publication of this paper. To analyze the test data via the provided software, we recommend using a system with at least 64 GB of RAM.

